# DNA Actively Regulates the “Safety-Belt” Dynamics of Condensin during Loop Extrusion

**DOI:** 10.1101/2025.04.29.651239

**Authors:** Jinyu Chen, Cibo Feng, Yong Wang, Xiakun Chu

**Affiliations:** Advanced Materials Thrust, Function Hub, The Hong Kong University of Science and Technology (Guangzhou), Guangzhou 511400, China; College of Life Sciences, Zhejiang University, Hangzhou, 310027, China; Guangzhou Municipal Key Laboratory of Materials Informatics, The Hong Kong University of Science and Technology (Guangzhou), Guangzhou 511400, China; Division of Life Science, The Hong Kong University of Science and Technology, Clear Water Bay, Hong Kong SAR 999077, China

## Abstract

Condensin, a protein complex of the SMC (Structural Maintenance of Chromosomes) family, is essential for genome folding through its active DNA loop extrusion activity. Condensin contains a binding interface between its Ycg1 (HEAT-repeat) and Brn1 (kleisin) subunits that acts like a “safety belt” to trap DNA and prevent its dissociation during loop extrusion. The entrapment of DNA within the binding pocket of the SMC complex is crucial for ATPase activity and the asymmetric loop extrusion. However, the molecular mechanism underlying DNA entrapment remains unclear, hindering our understanding of how condensin functions at a molecular level. Here, we employ multiscale molecular dynamics simulations to reveal how DNA actively modulates condensin’s safety-belt dynamics. Using all-atom simulations combined with AlphaFold3 predictions, we demonstrate that DNA binding markedly stabilizes the condensin safety belt, facilitating loop extrusion progression. Coarse-grained simulations capture the entire DNA entrapment process, highlighting an active regulatory role of DNA: DNA positioned outside the safety belt triggers its opening, whereas DNA within promotes closure, enhancing complex stability. Kinetic analyses further reveal that the rate-limiting step in DNA entrapment depends critically on the tightness of the safety belt. A loose safety belt makes the stable closure of its “latch” and “buckle” components rate-limiting, whereas a tighter safety belt shifts the barrier to initial DNA entry. Additionally, we find that latch-buckle interaction strength regulates condensin’s sliding motion along DNA. Our findings provide a comprehensive molecular framework illustrating how DNA actively guides condensin’s safety-belt dynamics, substantially advancing our mechanistic understanding of chromosome loop extrusion.

## Introduction

Mitosis and meiosis are fundamental processes of cell division. During these processes, the stable and orderly transmission of genetic information is crucial for maintaining species continuity and the healthy development of individuals [1–3]. The spatial organization of chromosomes is of paramount importance in mitosis, and achieving this organization relies on structural maintenance of chromosomes (SMC) complexes such as condensin.

Condensin, a five-subunit complex of the SMC family, plays a vital role in condensation and segregation during this process [4–6]. It is primarily composed of two key parts: the “SMC dimers” subcomplex and the “non-SMC” subcomplex [7]. The SMC dimers subcomplex consists of a heterodimer formed by SMC2 and SMC4 subunits and the non-SMC subcomplex composed of the kleisin and HEAT-repeat subunits [8]. Together, these two subcomplexes enable condensin to compact chromatin into highly ordered structures necessary for faithful chromosome segregation. Although condensin complexes were identified as pivotal players in mitotic chromosome formation, the the molecular mechanism by which condensin organizes chromosomes remains incompletely understood. Several models have been proposed over the years, and a simple and elegant mechanism that could explain most of the cellular tasks of SMC complexes called “loop extrusion” is one of the most widely accepted [4, 9–14].

Condensin complexes are characterized by conformational changes coupled with the ATP hydrolysis cycle and facilitate the lengthwise compaction of DNA through ATPase-driven loop extrusion [12, 15–20]. Current experimental evidence directly shows that the non-SMC subcomplex is responsible for DNA recognition and anchoring to chromatin, while the SMC2-SMC4 subcomplex on the other side functions as the molecular motor to perform motor action, enabling asymmetric one-sided loop extrusion [10, 21]. The observation that DNA binding by the non-SMC subcomplex of condensin decisively enhances the activity of SMC ATPase indicates a multistep mechanism involved in the loading of condensin onto chromosomes, which implies that the DNA binding site in the non-SMC subcomplex might be able to act as a sensor that triggers activation of the SMC2–SMC4 ATPase activitys [22]. The initial step in the condensin loading mechanism involves the binding of DNA to the non-SMC subcomplex, a process that is thought to occur independently of ATP hydrolysis [23]. A disruption in the interaction between DNA and the non-SMC subcomplex markedly affects the efficiency of ATP hydrolysis by the SMC2–SMC4 ATPase domains, highlighting that recognition of DNA by the non-SMC sub-complex is essential to activate the SMC2–SMC4 ATPases. [22].

A safety-belt mechanism, as revealed by DNA cocrystal structures, indicates that DNA is confined in a conserved, positively charged groove formed by Ycg1 HEAT-repeat and Brn1 kleisin subunits and is restricted by a ring-shaped Brn1 loop of non-SMC subcomplex (Figures 1A and 1B) [24]. This provides the structural basis for understanding how condensin anchors to chromosomes. DNA entrapment by the Brn1 loop and the Brn1 loop closure around the DNA double helix are essential to activate ATP hydrolysis by the SMC2-SMC4 ATPase domains [22, 24]. Although DNA can access the basic Ycg1–Brn1 groove without being encircled by the Brn1 loop, it is only through latch-mediated entrapment that the affinity is sufficiently increased to enable stable interaction with chromatin [24]. Since the “latch” and “buckle” segments can engage with each other even without DNA, the latch must temporarily disengage from the buckle to allow DNA to enter the Brn1 kleisin loop [24]. However, if the opening of the safety belt is a rare event, the efficiency of DNA entrapment would be exceedingly low. On the other hand, if the safety belt opens too readily, it could compromise the stability of ATP hydrolysis due to frequent and uncontrolled opening events. Therefore, understanding how the safety-belt mechanism achieves a balance between effective DNA entrapment and structural stability is essential for elucidating its functional mechanism.

**Figure 1.**
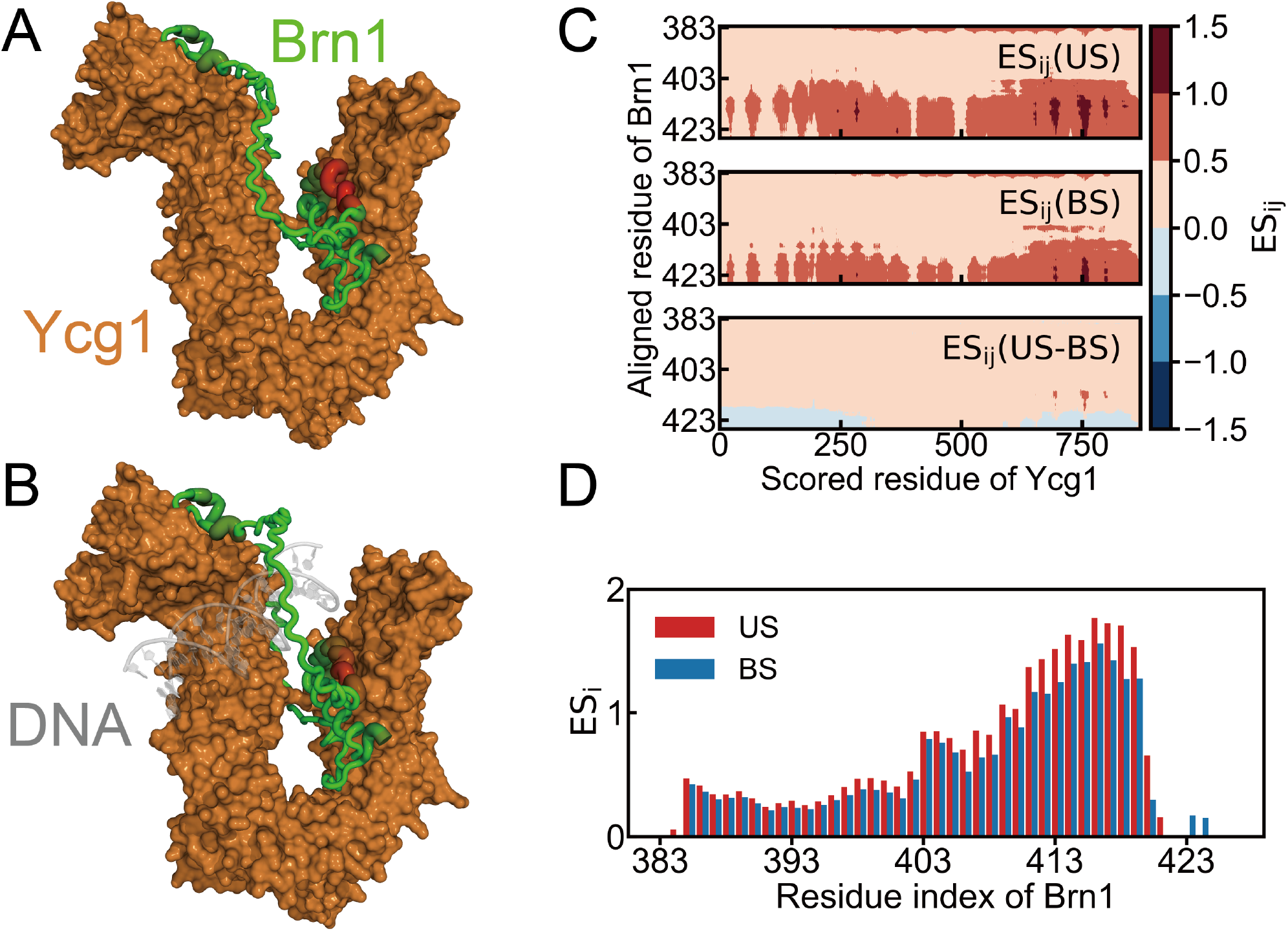
Effective strain results obtained from the AlphaFold3 predicted structure. The ES_*i*_ values are projected onto the corresponding prediction Brn1 kleisin structures of US (A) and BS (B). The thickness and color of the tubes in the predicted Brn1 kleisin structures denote the magnitudes of ES_*i*_, where a gradient from green to red represents increasing ES_*i*_ values. (C) ES_*i*_*j* matrix of the Brn1 loop highlights significant differences between the BS and US states, with Ycg1 as the scored residues and Brn1 as the aligned residues. (D) Local deformation per residue measured by ES*i* between the BS state and the US state in the Brn1 loop.

Previous studies have suggested, based on biochemical and structural evidence, that cohesin and condensin may topologically constrain DNA [25–31]. However, simultaneous closure of all three SMC–kleisin ring interfaces does not inhibit DNA loop extrusion by cohesin and condensin, thereby challenging the hypothesis that DNA passes through the SMC–kleisin ring in a strictly topological manner [11, 15]. In the hold-and-feed model, condensin is thought to load a DNA loop into the SMC–kleisin ring in a “pseudotopological” manner, whereby the DNA is topologically entrapped within two distinct kleisin chambers, referred to as the “motor” and “anchor” chambers. The topologically bound DNA segments within the safety belt during the SMC reaction cycle are thought to determine the directionality of DNA loop extrusion [15]. A recent study suggested that both cohesin and condensin are believed to extrude DNA loops in an asymmetric manner, and the frequent switching of extrusion direction has been proposed as a possible explanation for the experimentally observed symmetric loop extrusion [9]. Notably, unlocking the strict separation between the “motor” and “anchor” chambers converts condensin from a one-sided to a bidirectional DNA loop extruder. Therefore, a detailed understanding of the safety belt mechanism may provide important insights into the mechanistic differences in loop extrusion between condensin and cohesin.

Here, we use an integrated multiscale computational approach, combining AlphaFold3 prediction, all-atom molecular dynamics (MD) simulations, and coarse-grained (CG) modeling, to investigate the dynamics of condensin’s safety belt. We aim to elucidate the entire dynamic process of DNA entrapment by the safety belt and to identify several key intermediate states along this pathway. Our results reveal that DNA guides the modulation of the safety belt: it promotes safety belt closure when positioned inside and facilitates opening when outside, thereby ensuring efficient DNA entrapment and maintaining structural stability during ATPase activity. Furthermore, we demonstrate that the latch-buckle interaction strength is crucial for regulating the efficiency of DNA entrapment, revealing that a biologically optimized balance must exist to ensure both efficient DNA entrapment and stable ATPase activation. In addition, we found a strong correlation between the stability of the safety belt and the sliding of DNA. By uncovering how DNA modulates condensin’s safety-belt dynamics, our work provides crucial insights into the molecular basis of condensin’s function in genome organization.

## Results

### AlphaFold3 Predictions of Safety-Belt Flexibility in DNA Binding State (BS) and Unbinding State (US)

Recent advances in neural network-based protein structure prediction methods, such as AlphaFold [32], have enabled computational exploration of protein conformational diversity. Further studies have demonstrated that these predictive models can not only accurately reproduce experimentally determined static structures but also sample multiple conformational states by adjusting input parameters [33]. Additionally, prediction scores such as the predicted local distance difference test (pLDDT) and predicted aligned error (PAE) have been shown to strongly correlate with intrinsic protein conformational dynamics, allowing the identification of flexible regions [34]. Moreover, structural changes predicted by AlphaFold have been successfully applied to assess the impact of single-point mutations on protein conformation, with results showing significant agreement with experimental data [35]. These findings highlight the potential of predicted scores in AlphaFold to capture aspects of protein dynamics, providing new insights into conformational transitions and functional regulation.

Here, we first compared the behavior of the Brn1 kleisin loop in the DNA-bound (BS) and unbound (US) states within the safety belt. To analyze its dynamics and identify functionally relevant conformational changes, we assessed local deformation using the effective strain (ES) per residue, ES_*i*_, as a measure of flexibility. This method is similar to the local deformation concept proposed by McBride J. M. et al. [35], with the key difference being that we compute the average normalized PAE for each aligned residue *i* as the mean of PAE_*ij*_*/*r_*ij*_ (ES_*ij*_) for all scored residues *j* within 13 Å across all ten predicted structures. We extracted Brn1 kleisin loop residues as aligned residues and present the PAE matrix relative to the Ycg1 HEAT subunit, which serves as a more suitable reference due to its overall rigidity. To remove distance bias, we normalized the PAE by the distance between residue pairs in the corresponding predicted structure, enabling a fairer comparison of structural stability across regions and interaction scales. Higher normalized PAE values indicate that aligned residue *i* has greater uncertainty relative to scored residue *j*, while lower values indicate greater certainty. Consequently, a higher value of ES_*i*_ corresponds to increased flexibility in the local structural region, while a lower value of ES_*i*_ corresponds to decreased flexibility in the local structural region (see Methods for details). We projected ES_*i*_ directly onto the structure of the Brn1 kleisin loop to clearly visualize how the presence or absence of DNA affects the structural stability of the Brn1 kleisin loop (Figures 1A and 1B). Notably, in the presence of DNA, ES_*i*_ values are markedly reduced in a segment of the loop (∼residues 383-423), indicating enhanced local stability.

To further investigate the underlying cause of this decrease in ES_*i*_, we compared the ES_*ij*_ between the BS and US predicted structures in this region. As shown in Figure 1C, the ES_*i*_ shows a substantial overall reduction from the US to the BS state, clearly indicating the overall stabilizing effect of DNA in this region. The comparison of ES_*i*_ between the US and BS states in this region reveals more detailed insights (Figure 1D). This effect of stability is more pronounced in regions with higher normalized PAE values, while in regions with lower normalized PAE values (∼residues 383–403, *α*1 helix), the effect is relatively smaller but still exhibits an overall stable trend. The observed ES_*i*_ patterns are robust across a range of reasonable cutoff values, further supporting the reliability of this analysis (Figure S1). These findings suggest that DNA binding stabilizes specific flexible regions within the Brn1 loop, which is the main active region of the safety belt and is closely associated with the opening and closing motions between the latch and buckle. This stabilization may play a critical role in the closure of the Brn1 loop and its subsequent functional execution.

### Atomic Scale Insights into DNA-Stabilized Safety Belt Mechanism: Evidence from Unbiased MD and Steered MD Simulations

The prediction results of local deformations suggest that the DNA entrapment by the safety belt can facilitate the closure of the Brn1 loop. As AlphaFold3 primarily provides static structural predictions, additional validation through MD simulations or experimental approaches is required to assess the conformational dynamics. To further investigate how the safety belt maintains structural stability following DNA entrapment, we constructed two simulation systems, one with DNA and one without, and then performed unbiased all-atom MD simulations on each. The BS state was retrieved from the Protein Data Bank (PDB:5OQO). The US state was generated by deleting DNA from the PDB. The missing residues are reconstructed (see Methods for details). As shown in Figure 2A, the structural schematic illustrates key structural concepts within the complex, particularly the latch region, buckle region, and the primary active region (PAR). The pale green linker represents the missing portion of the Brn1 kleisin loop, while the Brn1 also contains four *α* helices (*α*1–*α*4). To quantitatively assess the structural dynamics of the Brn1 loop at the residue level in the presence and absence of DNA, we first compared the root mean square fluctuation (RMSF) between the BS and US states. The RMSF profile of the US state is generally larger than that of the BS state, indicating that the presence of DNA within the Brn1 loop enhances the overall structural stability of the safety belt (Figure 2B). Notably, residues near position 425 in the BS state show a modest increase in RMSF compared to the US state. This results from a higher concentration of negatively charged residues in that region, which, together with the negative charge of the DNA helix, weakens local interactions at the protein-DNA interface. Figure 2C illustrates that the primary contributor to RMSF stabilization is the direct interaction between DNA and the primary active region (PAR, residues 383–456) of the Brn1 loop. Moreover, the slightly higher contact number between the PAR and the rest of the protein (excluding DNA) in the BS state suggests that DNA binding may indirectly stabilize adjacent structural regions. These findings are consistent with the overall decrease in RMSF observed in the BS state.

**Figure 2.**
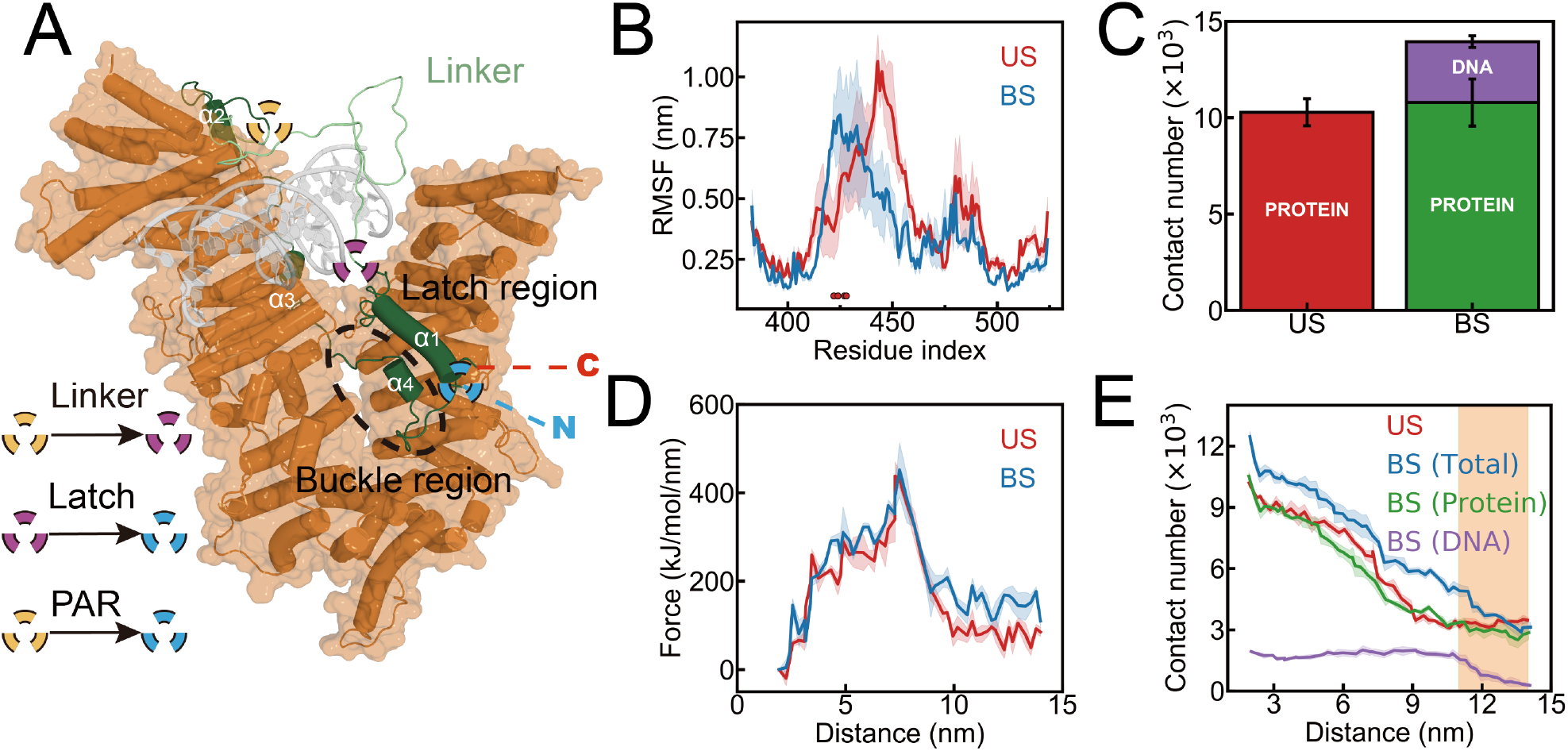
Stabilizing effect of DNA binding on the safety belt. (A) Structural schematic of the different regions in the protein complex. The subunit with the surface representation in orange indicating Ycg1 HEAT. The Brn1 kleisin is dark green. The linker region (pale green) is the missing region of Brn1 in crystal structure and reconstruction. Specific helices of Brn1 (*α*1-*α*4) are labeled for clarity. The C- and N-termini are marked with red and blue dashed lines, respectively. The latch and buckle regions are displayed in structure to distinguish functional regions. The symbols indicate linker, latch, and PAR elements as referenced in the legend. (B) RMSF of BS and US at the residue level. Negatively charged residues in regions where the BS value is higher than the RMSF value in the US state are marked with red circles. (C) Contact number between the primary active region and the remaining part in BS and US states during unbiased MD simulation. (D) Average force profiles as a function of distance obtained from steered MD simulations across three independent runs. (E) Contact number between the PAR and the remaining part in BS and US as a function of distance obtained from steered MD simulations across three independent runs. Here, in the BS state, the contact number between the PAR and DNA (purple), the PAR and the remaining protein (green), and the PAR and the total remaining part (blue) are shown. In the US state, the contact number between the primary active region and the total remaining part is represented in red.

The RMSF and contact number analyses from all-atom MD simulations showed that DNA stabilized the Brn1 loop by forming additional interactions. To further assess how this interaction influences latch disengagement from the buckle in the safety belt, we performed steered MD simulations. To preserve the flexibility of the buckle region at its interface with the latch region while preventing large-scale translations or rotations of the buckle region, positional restraints were applied to atoms of the Ycg1 subunit that do not directly interact with the PAR (see Figure S2 and Methods for details). We present the average force profiles as a function of the z-axis distance between the centers of mass (COMs) of the first C_α_ atom of the latch region of the Brn1 loop and Ycg1 subunits, derived from three independent steered MD simulations. Notably, the BS state exhibits generally higher force values compared to the US state (Figure 2D). The results indicate that DNA enhances the interaction network at the interface, leading to an increased force required for latch release. The presence of DNA stabilizes the interaction between the latch and the buckle, making the Brn1 loop less likely to open, which is important for the safety belt function and subsequent ATP hydrolysis. In the steered MD process, the latch region is pulled by a force applied along the z-direction at constant velocity, leading to its gradual detachment from the buckle region. The changes in the contact number between the PAR and the remaining part during this process are shown in Figure 2E. Even under steered MD, the comparison of contact numbers between the US and BS states remains highly correlated with the presence or absence of DNA near the crystal structure in the unbiased simulation (Figure 2C). The difference in contact number between US and BS comes mainly from the contact between DNA and the PAR. Quantitative analysis of contact numbers reveals comparable interaction profiles between the US and BS states when excluding contacts between DNA and the PAR of the Brn1 loop in the BS state (Figure 2E). In particular, the number of contacts between BS (total) and US reaches nearly the same value at almost the same time after the elimination of the interactions between the PAR and DNA (yellow highlight), strongly suggesting that these specific contacts are critical for maintaining the closure of the Brn1 loop. Combined evidence from unbiased MD simulations and steered simulations demonstrates that DNA entry stabilizes the closed conformation of the Brn1 loop. This mechanistic insight suggests that DNA-mediated loop stabilization can constitute an essential prerequisite for subsequent activation of SMC ATPase activity, ensuring the stability of the loop extrusion process.

### Dynamics of DNA Captured by the Safety Belt and Its Role in Modulating Safety Belt Dynamics

AlphaFold predictions and all-atom MD simulations suggest that DNA entry into the Brn1 loop enhances the stability of the safety belt. However, these findings are confined to conformations near the closed crystal structure of the Brn1 loop, leaving the initial mechanism of DNA entry unresolved. Notably, our simulations indicate that the transition of the safety belt between the open and closed states is rarely observed within several microseconds in allatom simulation (Figure S3), highlighting a limitation in capturing large-scale conformational transitions at atomic resolution. To overcome this problem and investigate the mechanism of DNA entrapment at spatio-temporal scales inaccessible with all-atom models, we developed a CG model of the Brn1–Ycg1–DNA complex. This model incorporates several key designed interactions to facilitate efficient exploration of the DNA capture process (Figure 3). In particular, a nonspecific interaction between DNA and the Ycg1 subunit is considered, which, in contrast to classical protein–DNA recognition, may enable condensin to access open chromatin regions, such as transcriptionally active genes or nucleosome-depleted sites [36, 37]. This nonspecific interaction facilitates DNA sliding along the protein complex and enables nonspecific captured by the safety belt. By ensuring the diffusion coefficients (*D*) are close in order of magnitude to the measured movement of cohesins and condensins moving diffusively along naked DNA in vitro (Figures 3A and S4), and the DNA bending angle (*θ*) observed in the crystal structure is reproduced with a longer DNA sequence (Figure 3B), we evaluated a range of interaction strengths between the Ycg1 HEAT-repeat subunit and DNA [6, 38–44]. The simulation closely matches experimental data in both *D* and *θ*, supporting the choice of *ε*_LB_ = 0.6 as the optimal interaction strength between DNA and Ycg1 (see Methods for details). The loop regions of Brn1 are largely missing, and the structured regions make few contacts with DNA, resulting in essentially no native contacts in the available structures. In our all-atom simulations with loop reconstruction, DNA interactions were primarily mediated by charged residues. Based on these observations, we focused on modeling Brn1–DNA interactions through electrostatics (Figure 3C). The strength of the interaction between the latch and the buckle (*ε*_LB_) is challenging to determine, as there are no direct experimental data available. To avoid artifacts caused by an inappropriate choice of interaction strength, we systematically explored a range of *ε*_LB_ values, allowing the transition of the safety belt from a nearly open to a fully closed state during the simulation (see Figures 3D, 3E and Methods for details).

**Figure 3.**
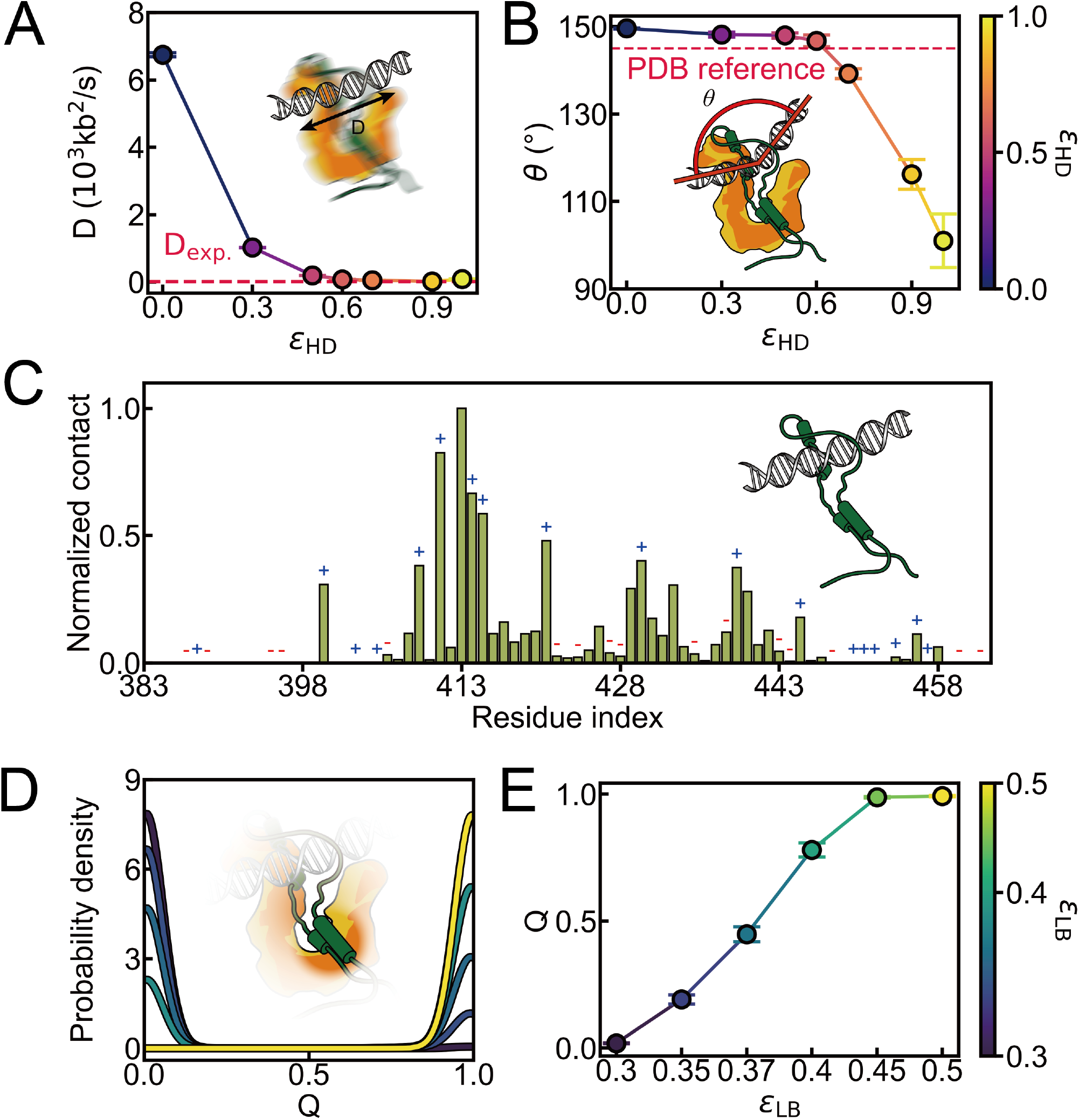
Analysis of the effects of different interaction parameters on various properties. (A) Diffusion coefficients of safety belt moving diffusively along naked DNA as a function of various values of *ε*_HD_ (0, 0.3, 0.5, 0.6, 0.7, 0.9, and 1), showing the impact of interaction between Ycg1 HEAT-repeat and DNA on the dynamics. The red dashed line is the mean experimentally accessible value of the *D* for cohesins and condensins diffusing along naked DNA. (B) Average bending angle of DNA as a function of *ε*_HD_, with a reference angle from the PDB (red dashed line). (C) Normalized contact between DNA and the Brn1 loop (residues 383-524), highlighting the variation in contact frequency across the residues. The blue “+” signs in the figure indicate positively charged residues, while the red “-” signs indicate negatively charged residues. (D) Probability density distributions of *Q* for different values of *ε*_LB_ (0.3 to 0.5). (E) Average *Q* as a function of *ε*_LB_, showing an increasing trend with the increasing value of *ε*_LB_.

Our simulations successfully recapitulated the complete process of DNA entrapment by the safety belt. To investigate whether this entrapment stabilizes the safety belt and to elucidate the role of DNA in modulating its dynamics, we employed the linking number to distinguish states whether DNA is topologically entrapped by the Brn1 loop (see Methods for details). Specifically, *S*_nl_ represents a state where DNA remains outside the loop, whereas *S*_l_ denotes a topologically entrapped state (Figure 4A). We then computed the average fraction of native contacts (*Q*) between the latch and buckle to quantify the extent of safety belt closure across trajectories. A higher *Q* value reflects more frequent closure, while a lower *Q* indicates a more frequent open conformation. Compared to the DNA-free state (*S*_f_), the *Q* value decreases in the *S*_nl_ state, but increases in the *S*_l_ state (Figure 4B). These results suggest that DNA can act bidirectionally: promoting opening when put outside the loop, and stabilizing closure upon entry. This DNA-dependent stabilization is consistent with previous all-atom MD simulations and AlphaFold3 predictions. The observed trends are robust across a broad range of *ε*_LB_ and *ε*_HD_ values (Figure 4B and Figure S5). Notably, when the latch–buckle interaction becomes excessively strong (e.g., *ε*_LB_ = 0.45), the modulatory effect of DNA is largely abolished, likely due to saturation of the closure interface. Interestingly, analysis of the DNA–Brn1 loop contacts reveals a compensatory mechanism: as *ε*_LB_ increases, a more stable loop conformation exposes additional DNA-binding sites, resulting in stronger interactions with DNA (Figure 4C). This is evident in the continuous increase in contact numbers in the *S*_nl_ state, while the *S*_l_ state shows only marginal changes. Intriguingly, these findings suggest that DNA, acting as a passive “passenger”, may guide its own entrapment by promoting tighter interactions when the safety belt adopts a more closed conformation.

**Figure 4.**
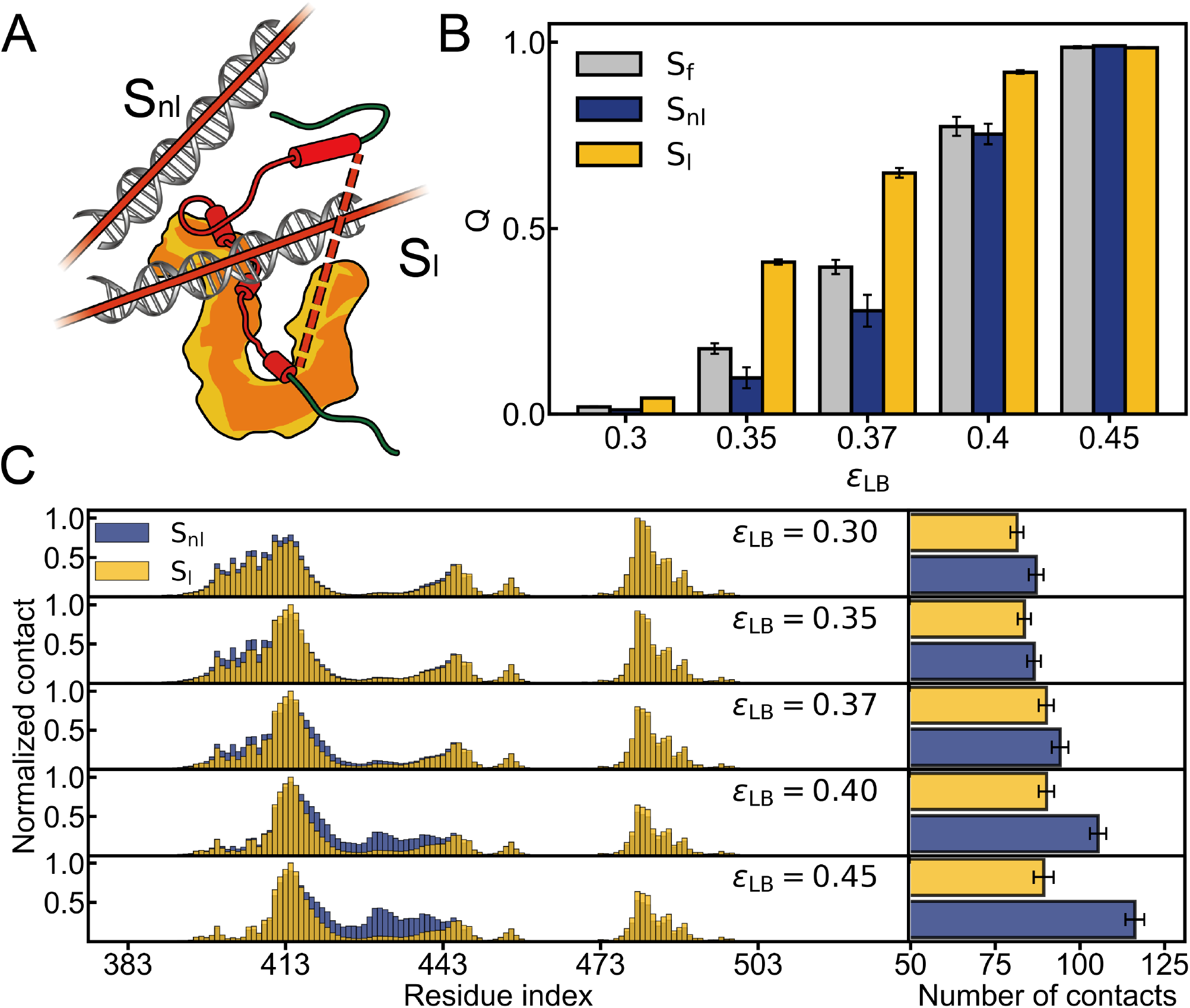
(A) Schematic illustration of *S*_nl_ and *S*_l_. The red regions represent the two loops constructed for calculating the linking number: the DNA is assumed to form an infinitely large closed loop, while the Brn1 loop together with the dashed segment forms the other closed loop. (B) Comparison of the fraction of native contacts in *S*_nl_ and *S*_l_ under different interaction strengths (*ε*_LB_). (C) Normalized number of contacts between the DNA and the Brn1 loop of the *S*_nl_ and *S*_l_ under the parameter of *ε*_LB_=0.3, 0.35, 0.37, 0.4 and 0.45. The number of contacts is normalized by dividing it by its maximum value. The average contact numbers of *S*_nl_ and *S*_l_ at the corresponding parameters are shown on the right.

### Effect of Latch-buckle Interaction Strength on the Kinetics of DNA Entrapment and Diffusion along DNA

It is now well established that DNA located outside the Brn1 loop promotes the opening of the safety belt, whereas DNA entrapped within the loop stabilizes its closure. This effect is modulated, at least in part, by the interaction strength between the latch and buckle, denoted as *ε*_LB_. To examine how variations in *ε*_LB_ influence the kinetics of DNA entrapment, we analyzed 50 independent CG simulation trajectories for each interaction strength. Throughout the simulations, we identified four different states: (i) DNA unbound to the Ycg1–Brn1 complex (US), (ii) DNA bound to the protein complex but located outside the Brn1 kleisin loop (IS_out_), (iii) DNA surrounded by the Brn1 kleisin loop but with an unclosed latch–buckle interface (IS_in_), and (iv) DNA fully entrapped by a closed loop (BS)

(Figure 5A). The state classification was based on a combination of the *Q*, the number of contacts between DNA and the Ycg1–Brn1 complex, and the topological linking number (Figure S6).

**Figure 5.**
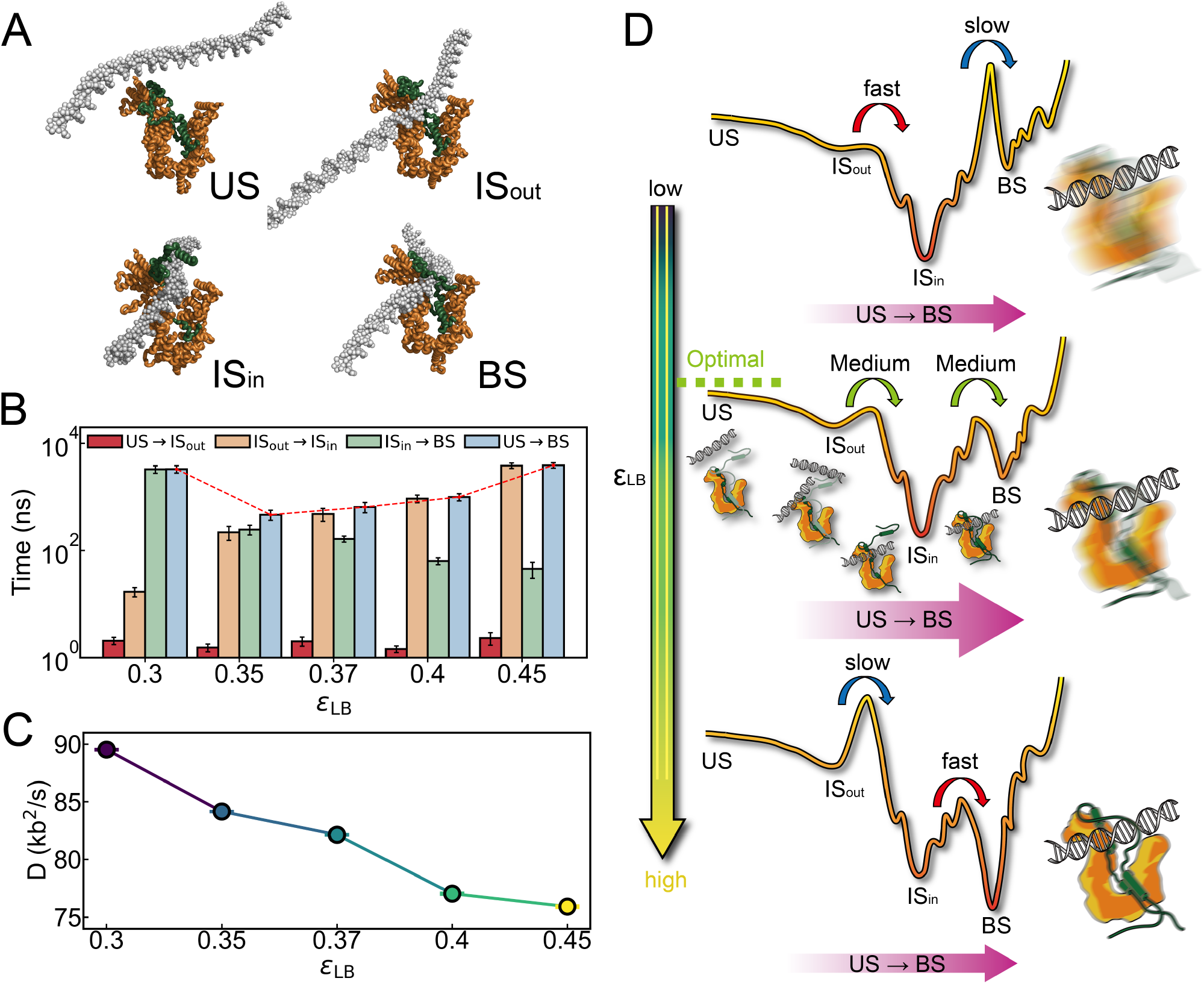
(A) Four binding states during DNA entry: US, IS_out_, IS_in_, and BS. (B) Transition times among the US, IS_out_, IS_in_ and BS, obtained from simulations in which DNA moves from outside the Ycg1-Brn1 complex into the binding pocket formed by the Brn1 loop and Ycg1 (the “safety belt”). The horizontal axis represents different values of the latch-buckle interaction strength *ε*_LB_, and the vertical axis indicates how each transition time (e.g., US →IS_out_, IS_out_ →IS_in_, IS_in_ →BS, US→ BS) depends on *ε*_LB_. (C) Dif fusion coefficients of safety belt moving diffusively along naked DNA as a function of various values of *ε*_LB_ (0.3, 0.35, 0.37, 0.4, 0.45), showing the impact of interaction between latch and buckle on the dynamics. (D) Scheme illustrating the primary transition pathways and approximate energy barriers involved in reaching the final bound state from the energy landscape perspective. The variation in the diffusion behavior of the safety belt as a function of *ε*_LB_ is illustrated on the right side of each case.

We subsequently measured the first passage times for the transitions US→ IS_out_, IS_out_→ IS_in_, IS_in_ →BS, and the complete transition US→ BS (Figures 5B and S7). Our results reveal that increasing *ε*_LB_ prolongs the transition from IS_out_ to IS_in_, suggesting that stronger latch–buckle interactions hinder loop opening and thus delay DNA entry. Conversely, the transition from IS_in_ to BS becomes markedly faster with increasing *ε*_LB_, reflecting facilitated loop closure once DNA is in place. The opposing influences of these two steps result in a non-monotonic transition time dependence of the process from US to BS under different *ε*_LB_, which initially decreases and subsequently increases. The *D* has been shown to influence condensin in locating appropriate anchoring sites and promoting linear compaction of chromatin [44]. Motivated by this, we hypothesized that the stability of the safety belt structure might similarly regulate the diffusion of DNA. To test this idea, we analyzed the relationship between the strength of latch–buckle interactions and the rate of one-dimensional linear diffusion of the safety belt along DNA. Our results revealed a clear inverse correlation: stronger interactions between the latch and buckle significantly slowed the diffusion of the safety belt on DNA (Figures 5C and S8).

A schematic free energy landscape illustrating these kinetic trends is shown in Figure 5D. Mechanistically, enhanced latch–buckle interactions impede DNA surrounded by the Brn1 loop (IS_out_ →IS_in_), but promote rapid closure of the Brn1 loop once DNA has entered (IS_in_ →BS). From a functional standpoint, strengthening ε_LB_ increases the stability of the fully engaged BS state, which in turn can facilitate the hydrolysis of ATP by the SMC2–SMC4 ATPase domains [24]. However, if ε_LB_ becomes too strong, the energetic barrier of DNA entrapment by Brn1 loop becomes prohibitively high, compromising the efficiency of DNA capture. Together, these findings suggest that an optimal latch–buckle interaction strength must balance structural stability with accessibility. The observed phenomena remain robust over a wide range (Figure S9). Moreover, the strength of the safety belt can also regulate DNA diffusion during the process of loop extrusion, thereby assisting condensin in locating appropriate anchoring sites and facilitating linear compaction of the chromatin [9, 44].

## Discussion

Through integrative multiscale computational approaches combining AlphaFold3 predictions, all-atom, and CG MD simulations, this study reveals the guiding roles of DNA in modulating the conformational dynamics of the safety belt, a critical DNA binding site of the condensin SMC complex anchoring to chromatin. Our results demonstrate that DNA binding not only stabilizes but also modulates the kinetic mechanism of the safety belt, thereby potentially regulating subsequent ATPase activity.

Using AlphaFold3 predictions, we observed that the presence of DNA significantly reduces local flexibility in the Brn1 loop. The effective strain analysis according to PAE score indicated that the BS state exhibits markedly lower flexibility compared to the US state. Specifically, the reduction in ES_*ij*_ and ES_*i*_ values beyond residue 400 suggests that DNA binding preferentially stabilizes the more flexible regions of the Brn1 loop. Notably, although the changes within the α1 helix region (residues 383–400) are more modest, a clear downward trend is still observed, suggesting that overall stability is indirectly influenced by DNA association. AlphaFold has demonstrated remarkable success in predicting static protein structures. While it remains an open question whether AlphaFold has fully captured protein dynamics, recent studies utilizing AlphaFold to predict conformational ensembles and to assess the impact of point mutations suggest that it may have implicitly learned key aspects of protein dynamics information [33, 35, 45]. Motivated by these observations, we explored the use of an effective strain metric derived from the PAE matrix to assess the impact of DNA binding on protein conformational changes. The results show strong agreement with both all-atom and CG MD simulations, supporting the potential of extracting dynamic insights from AlphaFold predictions.

Complementary to the AlphaFold3 predictions, unbiased MD simulations revealed that DNA binding significantly reduces the RMSF of the Brn1 loop (Figure 2B). The enhanced stability in the BS state was further supported by an increased contact number between the Brn1 loop and DNA, which indirectly stabilizes interactions with other protein regions (Figure 2C). In steered MD simulations, a higher force profile was observed to detach the latch from the buckle in the presence of DNA, indicating the stronger latch-buckle interaction induced by DNA binding (Figures 2D and 2E). Together, these results suggest that DNA plays an important role in maintaining the closed conformation of the safety belt, a state that is crucial for the subsequent activation of ATPase activity in the SMC complex.

To capture the large-scale dynamics of DNA entry into the safety belt, CG simulations were employed. We successfully observed the entire process of DNA entrapment by the safety belt. The simulations revealed the guiding functions for DNA: when located outside the Brn1 loop, it facilitates opening, whereas once inside, it promotes closure of the loop (Figure 4B). Four key states are identified: US, IS_out_, IS_in_ and BS, respectively (Figure 5A). In previous experiments, Kschonsak et al. observed a slight DNA upshift even under conditions where the Brn1 kleisin loop was covalently crosslinked [24]. We propose that this phenomenon may result from the IS_out_, in which the DNA does not enter the Brn1 loop but remains associated in close proximity to the safety belt structure. Although covalent closure of the loop prevents topological entrapment, the intrinsic flexibility of the kleisin linker may allow conformational rearrangements and expose the positively charged DNA-binding groove formed by the Ycg1–Brn1 subcomplex, thereby enabling DNA to associate through non-topological interactions. However, because this interaction lacks the topological constraint provided by loop entrapment, it is likely to be structurally unstable and easily disrupted by experimental conditions such as protein denaturation, mechanical stress, or electrophoretic separation. This may explain why the upshift is detectable but modest, occurring predominantly at higher protein concentrations. These findings collectively suggest that, beyond its role in topological entrapment, the safety belt may modulate non-topological DNA interactions through its conformational flexibility, offering structural insights into intermediate states during condensin loading. Notably, electrostatic interactions play a critical role in the positioning of DNA. Remarkably, even under conditions where ε_HD_ = 0, our model still shows a clear tendency for DNA to locate near the DNA-binding groove, highlighting the fundamental importance of electrostatics. This result is in strong agreement with a previous blind prediction that neglected Brn1 [46].

Then we quantified the transition time between these states. Systematic variation of the latch–buckle interaction strength ε_LB_ uncovered a nonmonotonic effect on the overall kinetics of DNA entrapment by safety belt. Stronger latch–buckle interactions stabilize the closed state, making it harder for DNA to enter initially, but they also speed up the final closure once DNA is inside. These conflicting effects indicate that an optimal interaction strength is needed to balance efficient DNA capture with the stability of the BS state.

Notably, purely electrostatic interactions between DNA and the condensin complex are insufficient to explain several key experimental observations. For example, condensin’s diffusion along DNA and the local DNA bending induced by the safety belt cannot be accounted for by longrange electrostatics alone [38, 44]. Therefore, introducing additional non-covalent interactions between DNA and the safety belt is essential for a more realistic and functional model. Given that condensin captures DNA through a non-specific sequence mechanism [24], we developed a modified Go-like interaction potential that enables nonspecific DNA recognition, supports one-dimensional diffusion consistent with the measured diffusive movement of cohesins and condensins along naked DNA in vitro, and accurately reproduces local DNA bending observed in crystal structures (Figures 3A and 3B). By tuning parameters within biologically relevant ranges, our model closely matches experimental behavior. Remarkably, at near-physiological protein concentrations, we observed the safety belt sliding along DNA, consistent with prior in vitro findings. This semi-diffusive sliding may facilitate chromosome folding by allowing condensin to explore local DNA configurations. Additionally, this sliding suggests that short or linear DNA fragments, unable to be topologically trapped, may escape the safety belt through diffusion, a behavior also reported in previous studies [10, 24].

Notably, we found a clear monotonic relationship between the stability of the safety belt and the degree of DNA sliding, suggesting a potential role in controlling its movement. Meanwhile, our simulations suggest that excessive sliding speed can disrupt stable DNA entrapment, impairing safety-belt closure (Figure S10). This delicate balance between mobility and stability may act as a key regulatory mechanism in condensin’s function during chromosomal loop extrusion. Interestingly, although yeast condensin is considered an asymmetric and unidirectional loop extruder-reeling in DNA loops from only one side-experiments have shown that it transitions from a static to a sliding behavior under near-physiological ionic conditions [10]. Although salt concentration was not explicitly investigated, our preceding results imply that it may influence DNA-protein interactions and, consequently, the sliding behavior of DNA, which could play a role in regulating safety belt opening and closure.

Our study offers significant mechanistic insights, yet several questions remain. For example, the energetic contributions of specific residue interactions within the Brn1 loop and potential allosteric effects on other SMC subunits require further exploration. Recent evidence highlights asymmetric DNA loop extrusion as a shared mechanism among eukaryotic SMC complexes, supported by single-molecule and structural studies of condensin, cohesin, and SMC5/6 [9]. Notably, a safety belt–like structure has been identified in the Scc3–Scc1 subcomplex of the budding yeast cohesin complex [47], suggesting that, despite variations in composition and regulation across species, asymmetric loop extrusion may be a conserved feature of SMC complexes. Extending our model to other SMC complexes could reveal whether DNA-mediated safety belt regulation is a universal mechanism in SMC-mediated chromosome organization.

In summary, our findings demonstrate that DNA modulates the dynamics of the safety belt in the condensin complex actively. The functional cycle of this complex relies on a critical balance between the strength of latch–buckle interactions and conformational changes induced by DNA. These insights could have important implications for understanding the molecular basis of chromosome architecture and dynamics, and how SMC complexes like condensin orchestrate large-scale genome organization. Our multiscale simulation approach establishes a framework that can be extended to study other SMC complexes, bridging static structures and dynamic function.

## Methods

### All-atom MD Simulation

In this study, the initial structure of the all-atom simulation is constructed based on the crystal structure (PDB code: 5OQO) [24]. Missing residues are reconstructed using Modeller 10.4 [48]. The DNA-free structure is obtained by removing the DNA from the reconstructed structure. All systems are built using the web-based CHARMM-GUI Membrane Builder [49].Charmm36m was used to describe the interactions [50].The steered MD was conducted to compare the impact of the Brn1 loop encircling DNA on the safety belt. A single reaction coordinate was defined between two groups, the first C_α_ atom of the latch region in the Brn1 loop (the pulling group) and the Ycg1 (the reference group). An external harmonic potential was applied along the z-axis using the “direction” geometry, with pulling dimensions set to only affect the z-coordinate. The pulling was performed at a constant rate of 0.005 nm ps^−1^, with a force constant of 600 kJ mol^−1^ nm^−2^. The initial distance between the COMs of the two groups was used as a starting point, and the periodic boundary conditions were accounted for by updating the reference coordinates at each step. The protein is rotated in a way that the DNA binding groove is located in the direction of the z-axis. The atom positions of the protein with z *<* 14 nm in our box are fixed to ensure that the position of the Ycg1 HEAT-repeat does not change, and the remaining parts are not fixed to keep the binding position as flexible as possible (Figure S2). The steered MD runs of 2500 ps fully extract the latch region from the buckle region.

Unbiased MD and steered MD simulations were conducted using the GROMACS 2022.5 package with an integration time step of 2 fs. Coulomb interactions were shifted to zero between 1.0 and 1.2 nm, and a cutoff radius of 1.2 nm was applied for Lennard-Jones interactions. The system temperature was maintained at 310 K using a velocity-rescaling thermostat, and the pressure was kept at 1 bar in all directions with a Parrinello-Rahman barostat. Simulation boxes measured 17 ×17 ×17 nm^3^, which was filled with TIP3P water molecules, and 0.15 M NaCl was added to mimic the physiological condition. We performed three independent simulations for each system, each running for 1 *µ*s.

### Modelling the CG Structure of Brn1-Ycg1 in Complex with DNA

To investigate the large-scale dynamics of DNA entering the safety belt, we employed CG simulations. The initial structure for the simulation was constructed from the aforementioned all-atom Brn1-Ycg1-DNA complex with missing residues reconstructed. For the Brn1-Ycg1 complex, a residue-level CG model (AICG2+) was used, with each bead positioned at the C_*α*_ atom location [51]. Considering the lack of structural information for the missing region (residues 413–456) in Brn1, we removed the structure-based local contact potentials and retained only the generic local potentials derived from a “loop-segment library” based on Protein Data Bank statistics to better capture its dynamic behavior. These generic local potentials effectively describe the conformational distribution of unfolded states or intrinsically disordered regions by accounting for secondary structure propensities [52]. For DNA, we employed the 3SPN.2C model [53], which captures base-pairing, hybridization, groove widths, and local curvature. In this model, each nucleotide is represented by three beads corresponding to the phosphate, sugar, and base groups. To prevent the false capture of DNA, where one end of short linear DNA fragments may pass through the safety belt, we extended the original 18-bp DNA double helix by 40 bp at each end, resulting in a 98-bp DNA double helix structure. A spherical flat-bottom potential, with a radius of 10 nm, was applied between the center of mass of the original 18-bp DNA residues and the protein center. No periodic boundary conditions were used in any direction.

The Brn1-Ycg1 complex and DNA interact through long-range electrostatics, excluded volume, and a nonspecific interaction of the protein-DNA sequence, which expands the Go-like potential for the Ycg1-DNA interaction observed in the crystal structure. We first extract the native contact list between Ycg1 and DNA. These interactions were formed between a set of protein residues and related phosphate or sugar beads of DNA in the crystal structure. We then expanded this list by increasing or decreasing the index of phosphate and sugar in DNA to cover the entire 98-bp range of DNA. The index of Ycg1 in the list did not change, forming a Ycg1-protein interaction model without specific binding.

To determine the interaction strength between the DNA duplex and Ycg1, we first compared the mean experimentally accessible value of the *D* for cohesins and condensins diffusing along naked DNA. The experimentally measured range of D in vitro is approximately from 0.01 to 35 kb^2^*/*s. When *ε*_LB_ ≥ 0.5, the simulated diffusion coefficient D reaches values ranging from several tens to over a hundred kb^2^*/*s, approaching the upper bound of in vitro experimental measurements. Given that CG simulations generally overestimate diffusion due to reduced frictional damping [54], this level of agreement is remarkably close to experimental observations. Notably, when *ε*_LB_ ≤ 0.6, the *θ* observed in the simulations aligns well with that seen in crystal structures with a longer DNA fragment (PDB: 7Q2Z) [38]. By comparing the *D* and *θ* at different interaction strengths with the angle in the crystal structure, we determined that the interaction strength is 0.6 times the default value. Additionally, we focused on the impact of the interaction strength between the latch and the buckle in the safety belt model on the overall dynamics of DNA entry. To explore this, we tested a range of parameters with varying interaction strengths, specifically between 0.3 and 0.5, performing 10 independent simulations for each parameter. Each simulation system was run for a total of 100 *µ*s. The probability distribution of *Q* under these parameters is shown in Figure 3D. Our results demonstrate that the interaction strength between latch and buckle produces a shift in the mean value of *Q* from approximately 0 to 1 (Figure 3E), indicating that the safety belt can transition from almost not closed to fully closed under the tested conditions.

Our all-atom MD simulations reveal that the interaction between DNA and Brn1 is primarily electrostatic near the crystal structure (Figure 3C). Considering that there are only a few amino acid residues on Brn1 kleisin that have native contact with DNA in the crystal structure, and these residues are predominantly charged, we focus on electrostatic interactions and volume repulsion between DNA and Brn1 without adding additional Go-like potentia. Following the suggested parameters [53, 55], we assigned a charge of -0.6e to each phosphate bead for DNA-DNA electrostatic interactions, ensuring the appropriate DNA persistence length and accounting for the Oosawa-Manning condensation of counterions surrounding DNA. For Ycg1-Brn1-DNA electrostatics, we set the phosphate charges to -1e, as described in Refs [56, 57]. Electrostatics is modeled according to the Debye-Hückel theory.

All CG MD simulations were performed using GENESIS-2.1.2 with an integration time step of 0.01 ps [58]. Langevin dynamics with default parameters were employed at a temperature of 310 K. To investigate the kinetics of DNA entry under different conditions, 50 independent 10 *µ*s simulations were conducted for each condition. A summary of the simulation systems and durations is provided in Table S1. The total simulation time for the Brn1-Ycg1-DNA complex system was 6700 *µ*s, while for the Brn1-Ycg1 complex system it was 600 *µ*s. The ionic strength was set to 0.15 M.

### Data Analysis

AlphaFold is capable of predicting structural change and information about protein dynamics [33–35]. To quantify local deformation, we use a variable ES_*i*_, which is a sensitive measure of local structural change. The ES_*i*_ was defined as the mean relative change in inter-residue predicted aligned error (PAE) around a residue. For each residue *i*, we define a neighborhood *N*_*i*_ comprising *n*_*i*_ = |*N*_*i*_|, residues *j* ∈ *N*_*i*_ with *C*_α_ distances **r**_*ij*_ within 13 Å . The scenarios with and without DNA were constructed for comparison, and 10 sets of predictions were performed for each system. The ES_*i*_ is the average relative change in ES_*ij*_ over the *n*_*i*_ neighbors:

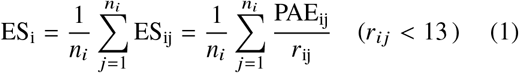

Visualization of the data was performed with Visual Molecular Dynamics (VMD) and Pymol software. Several GROMACS tools were used to analyze MD trajectories.

To examine how entry of DNA effects the stability of the safety belt, *gmx*_*mindist* was adopted with a cutoff distance of 0.6 nm to calculate the contact number. The interactions between the DNA and protein residues were described by the normalized number of contacts, which is defined by the ratio of the average number of contacts per frame of each residue to the maximum calculated above. RMSF was calculated with *gmx*_*rmsf* . The *Q* formed by the latch and buckle in the crystal structure is determined by the switching function:

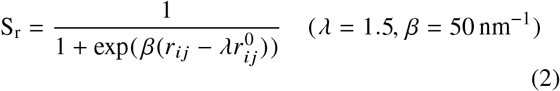

The probability distribution of *Q* was calculated by HISTOGRAM and was carried out with the aid of the PLUMED plugin (version 2.9.0) [59, 60].

To quantify the one-dimensional diffusion of the condensin along DNA, we calculated the mean square displacement (MSD) based on the index of the DNA base pair nearest to the binding site at each frame. Specifically, the MSD was computed as:

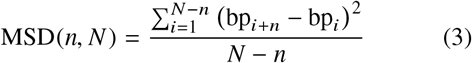

where bp_*i*_ denotes the base pair index closest to the protein binding site at frame *i, N* is the total number of frames in the trajectory, *n* is the frame lag, and Δ*t* is the time interval between two consecutive position measurements. Although this coordinate system is discrete, it captures the stepwise transitions of the protein binding site along the DNA sequence and is suitable for analyzing sliding or hopping behavior[61].

To estimate the diffusion coefficient *D*, we focused on the initial linear regime of the MSD curve, where MSD increases approximately linearly with lag time:

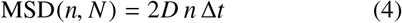

To ensure reliable fitting and minimize statistical uncertainty, we imposed a cutoff lag *n*_*c*_, chosen so that the relative error in MSD remained below 20% [61, 62]. The MSD variance was estimated as:

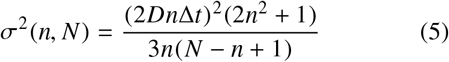

so that the cutoff was determined by satisfying:

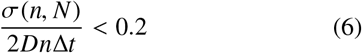

To minimize potential artifacts in the MSD analysis introduced by the flat-bottom potential (10 nm radius) between the COMs of the original 18-bp DNA residues and the protein complex, we excluded trajectory segments where the protein approached the ends of the DNA duplex. The DNA molecule spans 98 base pairs, with an average spacing of 0.34 nm per base pair. Thus, the central region of approximately 58 base pairs corresponds to a 20 nm potential-free zone (diameter, calculated as 10 nm radius × 2). To avoid edge effects, we only analyzed trajectory segments where the DNA base pair closest to the protein binding site continuously resided within the central 50 base pairs (well within the ∼58 bp potential-free zone) for at least 1000 frames. Based on this criterion, the number of valid trajectory segments obtained under different (ε_HD_) was 3 (ε_HD_ = 0), 182 (ε_HD_ = 0.3), 75 (ε_HD_ = 0.5), 54 (ε_HD_ = 0.6), 32 (ε_HD_ = 0.7), 55 (ε_HD_ = 0.9), and 11 (ε_HD_ = 1). For the latch-buckle interaction strength (ε_LB_), due to the availability of more extensive trajectories, we applied a stricter criterion, requiring the closest base pair to remain within the central 50 base pairs for at least 5000 frames. The resulting numbers of valid trajectory segments were: 114 (ε_LB_ = 0.3), 116 (ε_LB_ = 0.35), 108 (ε_LB_ = 0.37), 121 (ε_LB_ = 0.4), and 109 (ε_LB_ = 0.45).

Under our simulation conditions, we selected *n*_*c*_≈ 60 frames (6 ns) for segments of 1000 frames and *n*_*c*_≈ 300 frames (30 ns) for segments of 5000 frames. This choice balances statistical accuracy and time resolution and ensures that the estimated *D* reflect genuine sequence-position dynamics.

The bending angle of DNA is defined by the maximum angle of curvature formed by the line connecting the centroids of its corresponding base pairs.

The topological state between the Brn1 loop and DNA was quantified using the concept of “linking number”, a fundamental invariant in knot theory that measures the entanglement of two closed curves in three-dimensional space. For two non-intersecting closed curves *γ*_1_ and *γ*_2_, the linking number *Φ*(*γ*_1_, *γ*_2_), defined via the Gauss linking integral,

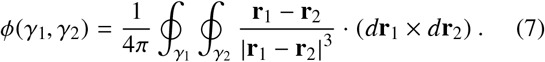

is an integer that remains invariant under continuous deformations, representing the number of times one curve winds around the other. Here **r**_1_ and **r**_2_ are the position vectors of points in the curve *γ*_1_ and *γ*_2_, respectively. In our molecular dynamics system, we modeled the Brn1 loop as a closed curve by connecting its termini with a straight line. This approximation is physically reasonable, as it captures the effective linking when the DNA is entangled by the Brn1 loop and the auxiliary segment. To treat the finite-length DNA duplex as a topologically closed curve, we extended the nearly linear DNA infinitely from its termini along the axial direction. Because contributions to the linking integral from distant segments are negligible, we truncated the extension beyond 4 nm from each terminus. The resulting DNA segment is sufficiently long relative to the Brn1 loop to effectively approximate an infinite-length DNA. The linking number between the Brn1 loop and the extended DNA segment was computed numerically. Convergence tests confirmed that extending the DNA beyond 106 nm did not alter the integral value, indicating a stable topological classification. Instances where the extended DNA overlapped with the Brn1 loop (resulting in non-integer values due to numerical singularities) were rare and excluded from analysis. The computed linking number was strictly quantized: *Φ*(*γ*_1_, *γ*_2_) = 0 indicated that the DNA duplex was located outside the loop without threading, whereas |*Φ*(**γ**_1_, **γ**_2_)| = 1 indicated that the DNA duplex passed through the Brn1 loop. This approach provided a robust metric to statistically resolve DNA-loop threading events in our simulations.

## Data availability

Data supporting the findings of this work are available in the main text, Methods, Supplementary Information, or Supplementary Data. Source data are provided with this paper.

## Acknowledgements

X.C. thanks Prof. Qianyuan Tang for useful discussion. X.C. acknowledges support from the National Natural Science Foundation of China (Grant Nos. 32201020 and 12474201), the General Program of the Guangdong Basic and Applied Basic Research Foundation (Grant No. 2024A1515010862), the Guangdong Scientific Research Platform and Projects for Higher-educational Institutions from the Department of Education of Guangdong Province (Grant No. 2023KTSCX169), and the Guangdong Provincial Project (Grant No. 2023QN10X037).

