## Supplementary material for "DNA Actively Regulates the “Safety-Belt” Dynamics of Condensin during Loop Extrusion": SI Text

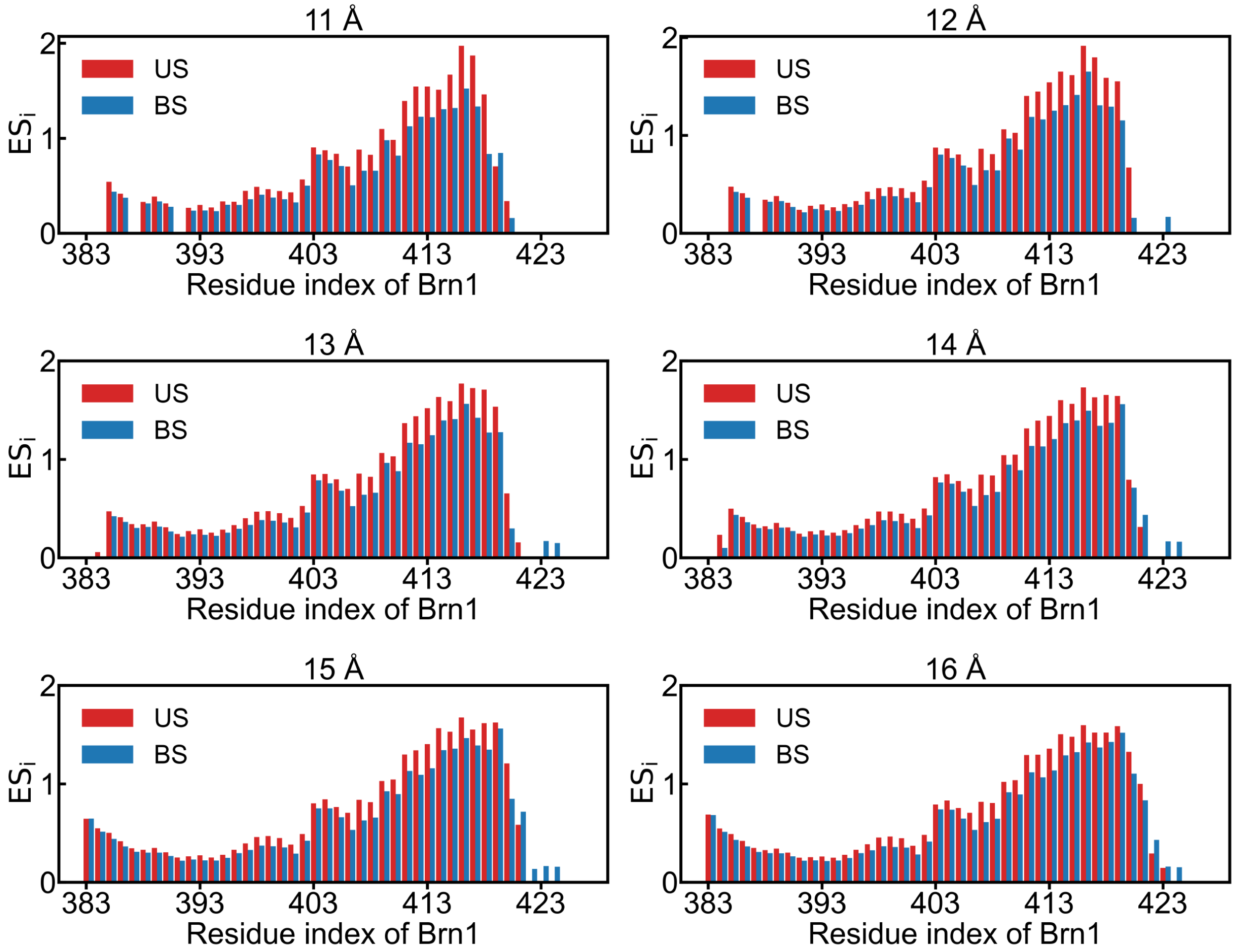


Figure S1: Local deformation per residue measured by $ES_i$ between the BS state and the US state in different distance cutoffs.

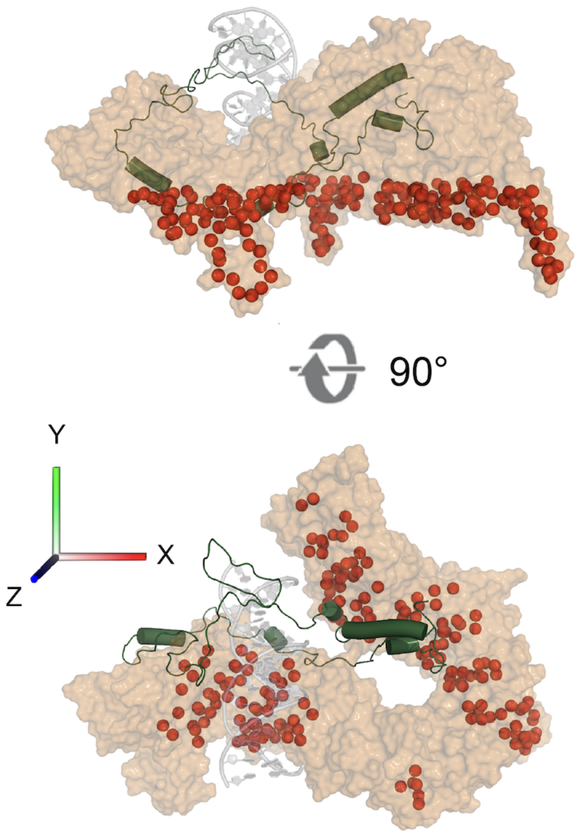


Figure S2: Constraints on the protein in the presence and absence of DNA. (A) System with DNA, where the red regions represent the $C_\alpha$ atoms subjected to restraints. (B) System without DNA, highlighting the same set of restrained $C_\alpha$ atoms in red.

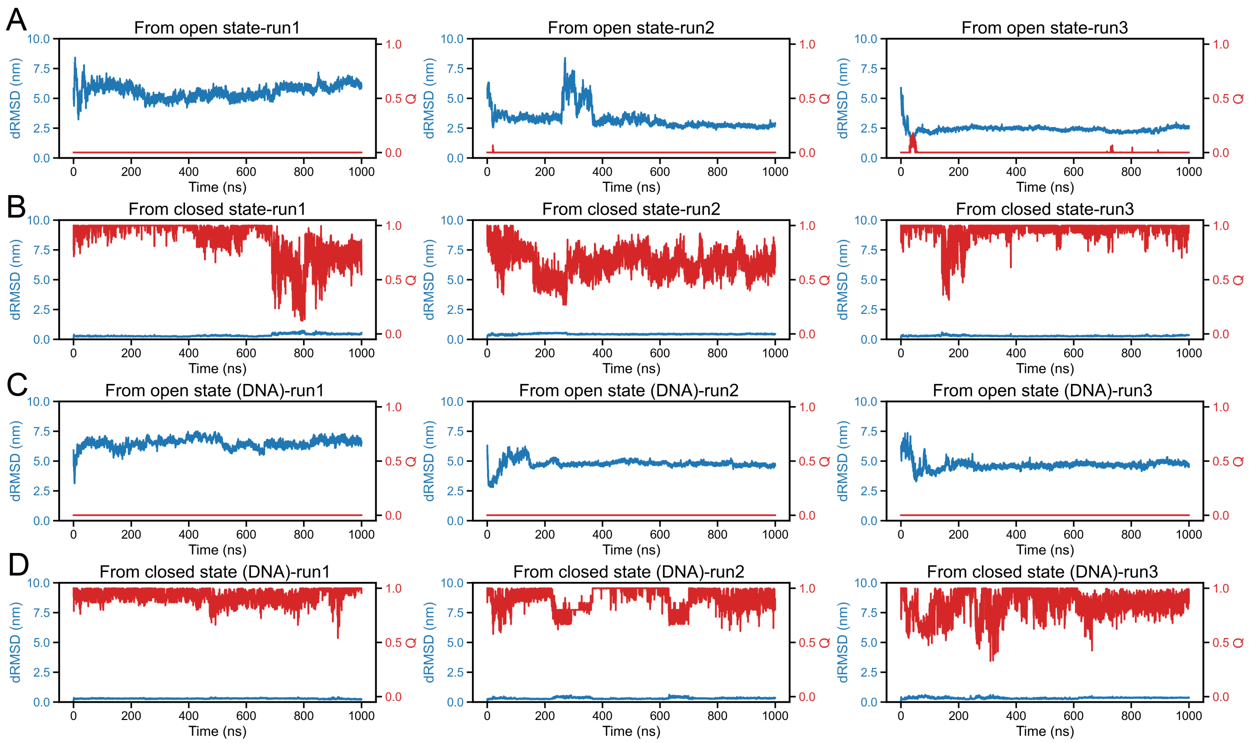


Figure S3: Time evolution of dRMSD and $Q$ from all-atom simulations under four conditions: (A) safety belt open, DNA absent; (B) safety belt closed, DNA absent; (C) safety belt open, DNA present; (D) safety belt closed, DNA present.

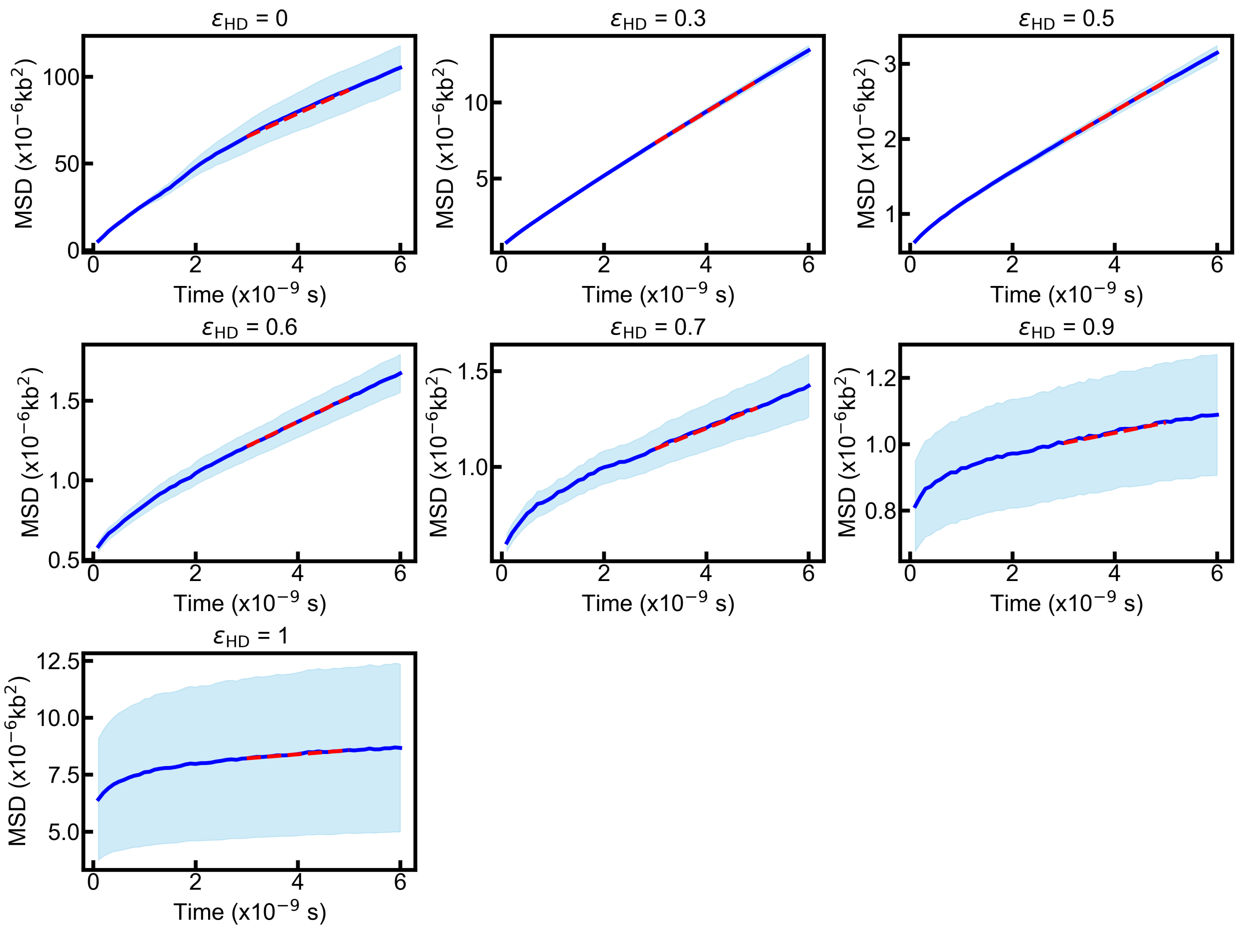


Figure S4: Mean square displacement (MSD) analysis of sliding motions of the safety belt on DNA as a function of time in different $\varepsilon_{\text{HD}}$. The mean MSD and standard error for the dataset are shown as a thick continuous line and shaded area, respectively. Diffusion coefficients were obtained from the slope of the red dashed line, which is a linear fit to the MSD curve within the specified time window.

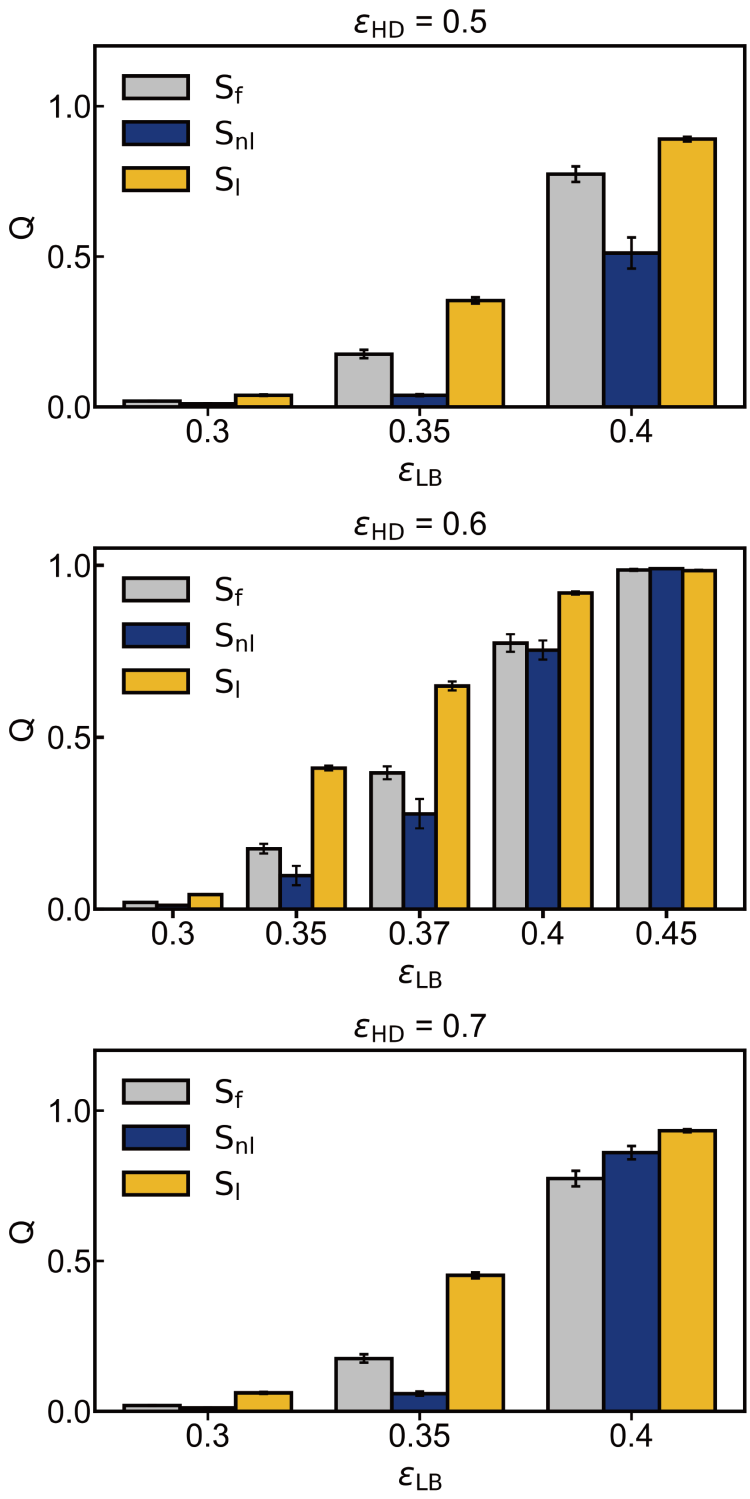


Figure S5: Effect of $\varepsilon_{\mathrm{LB}}$ on $Q$ between latch and buckle under different values of $\varepsilon_{\mathrm{HD}}$ (under $\varepsilon_{\mathrm{LB}} = 0.5$ (A), $\varepsilon_{\mathrm{LB}} = 0.6$ (B), and $\varepsilon_{\mathrm{LB}} = 0.7$ (C)). The values of $Q$ are shown for different states: $S_{\mathrm{f}}$, $S_{\mathrm{nl}}$, and $S_{\mathrm{l}}$ as a function of $\varepsilon_{\mathrm{LB}}$.

<table>
<tr><th>Linking number</th><th>Q</th><th>HEAT-DNA contact</th><th>Kleisin-DNA contact</th><th>state</th></tr>
<tr><td>--</td><td>--</td><td><0.5</td><td><0.5</td><td>US 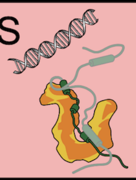</td></tr>
<tr><td rowspan="2"><0.8</td><td rowspan="2">--</td><td>>0.5</td><td>--</td><td rowspan="2">IS<sub>out</sub> 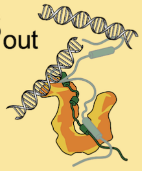</td></tr>
<tr><td>--</td><td>>0.5</td></tr>
<tr><td>>0.8</td><td>--</td><td>--</td><td>--</td><td>IS<sub>in</sub> 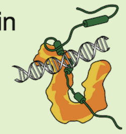</td></tr>
<tr><td>>0.8</td><td>>0.8</td><td>--</td><td>--</td><td>BS 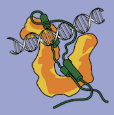</td></tr>
</table>

Figure S6: A table summarizing the different states based on the linking number, $Q$, DNA-HEAT contact, and DNA-kleisin contact. The states are classified as US (unbound state), $\text{IS}_{\text{out}}$ (intermediate state, outside), $\text{IS}_{\text{in}}$ (intermediate state, inside), and BS (bound state). The corresponding structures are displayed on the right.

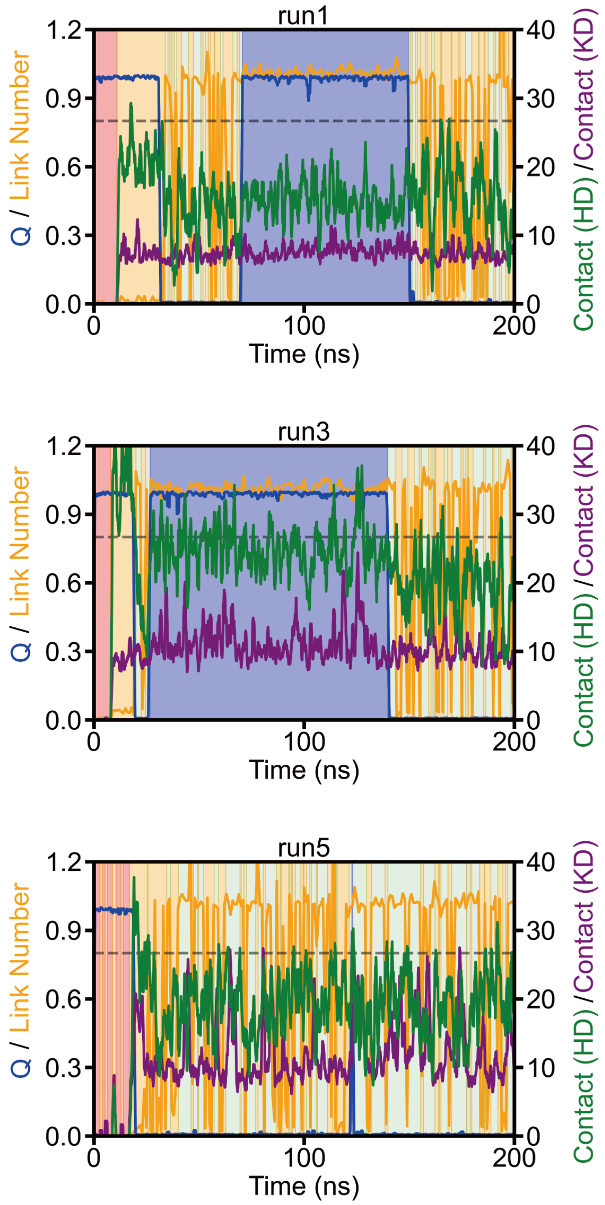


Figure S7: Representative trajectories reveal conformational transitions among four states under the values of $\varepsilon_{\mathrm{LB}} = 0.35$. Plots show the time evolution of $Q$, linking number, HEAT–DNA contacts (HD), and kleisin–DNA contacts (KD). Shaded regions indicate different states: red (US), orange ($\mathrm{IS_{out}}$), green ($\mathrm{IS_{in}}$), and purple (BS). The grey dashed line marks a reference value of 0.8 on the left y-axis used for linking number and $Q$.

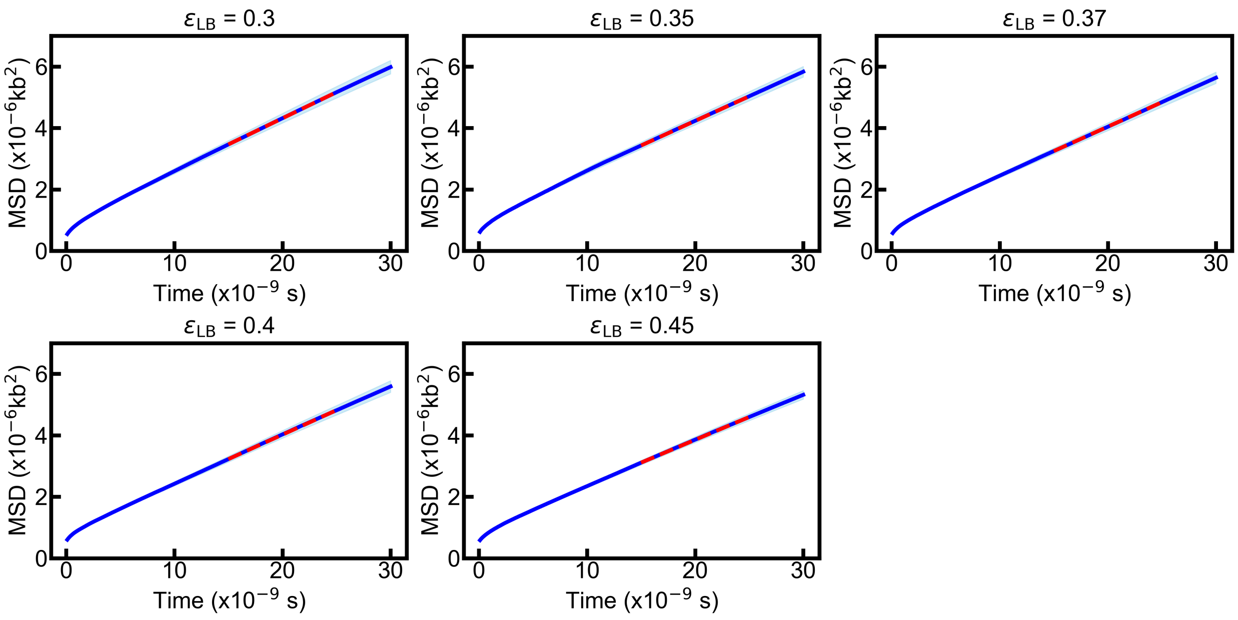


Figure S8: Mean square displacement (MSD) analysis of sliding motions of the safety belt on DNA as a function of time in different $\varepsilon_{LB}$. The mean MSD and standard error for the dataset are shown as a thick continuous line and shaded area, respectively. Diffusion coefficients were obtained from the slope of the red dashed line, which is a linear fit to the MSD curve within the specified time window. Same as Figure S4, but for different $\varepsilon_{LB}$.

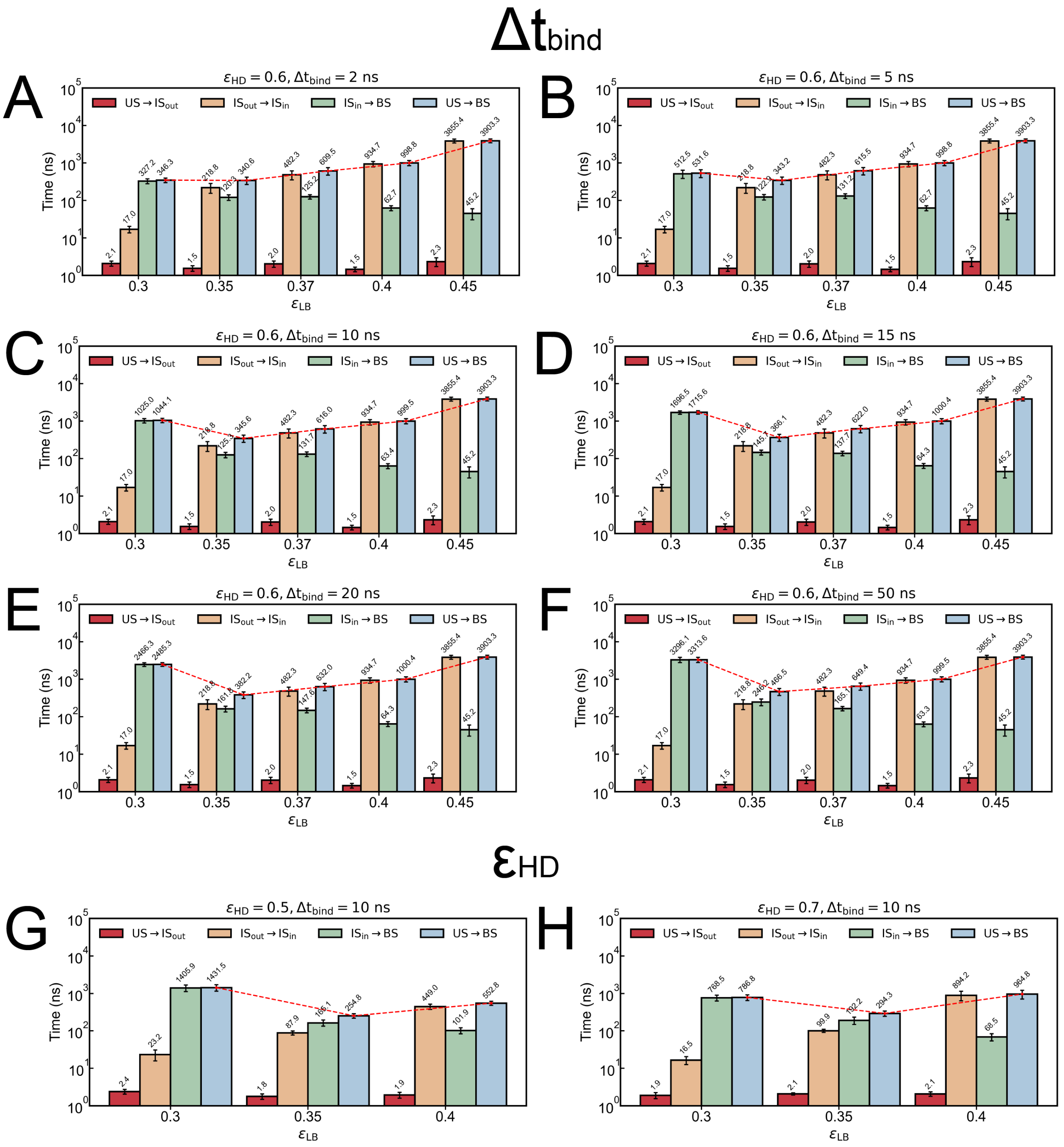


Figure S9: Transition time between different states under varying simulation parameters and conditions. (A-F) Results for different $\Delta t_{\text{bind}}$ values: (A) 2 ns, (B) 5 ns, (C) 10 ns, (D) 15 ns, (E) 20 ns, and (F) 50 ns, with $\varepsilon_{\text{HD}} = 0.6$, where $\Delta t$ defines the minimum duration a configuration must persist to be considered the BS state. The transitions include US $\rightarrow$ IS$_{\text{out}}$, IS$_{\text{out}}$ $\rightarrow$ IS$_{\text{in}}$, IS$_{\text{in}}$ $\rightarrow$ BS, and US $\rightarrow$ BS. (G-H) Results for different $\varepsilon_{\text{HD}}$ values: (G) $\varepsilon_{\text{HD}} = 0.5$ with $\Delta t_{\text{bind}} = 10$ ns, and (H) $\varepsilon_{\text{HD}} = 0.7$ with $\Delta t_{\text{bind}} = 10$ ns. The red dashed line shows the trend of transition time (US $\rightarrow$ BS) as a function of $\varepsilon_{\text{LB}}$.

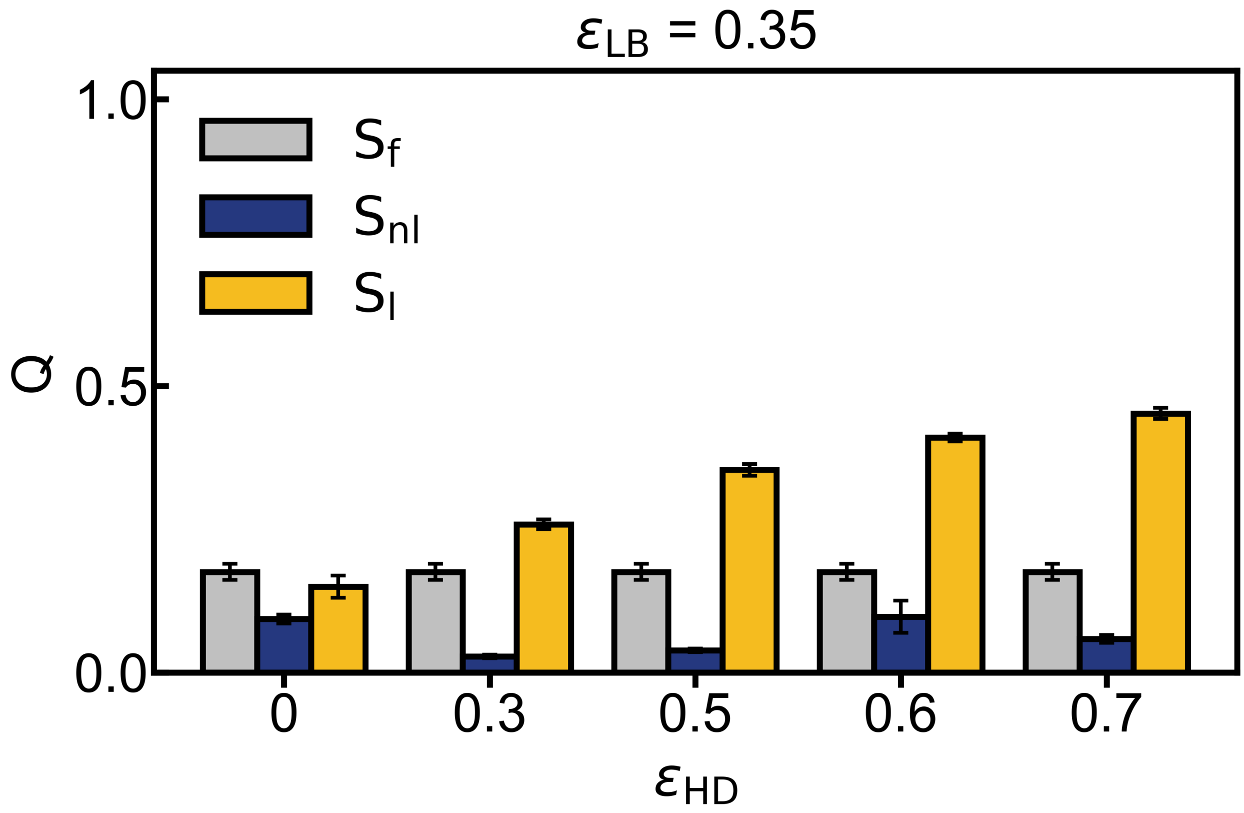


Figure S10: Effect of $\varepsilon_{\mathrm{HD}}$ on $Q$ between latch and buckle under the values of $\varepsilon_{\mathrm{LB}} = 0.35$. The values of $Q$ are shown for different states: $S_{\mathrm{f}}$, $S_{\mathrm{nl}}$, and $S_{\mathrm{l}}$ as a function of $\varepsilon_{\mathrm{HD}}$.

Table S1: Overviews of the simulation trajectories

| **Unbiased MD Simulation** | | | |
|---|---|---|---|
| **System** | **state** | **Duration** | |
| Brn1 + Ycg1 | | | |
| 1 | closed | $3 \times 1$ $\mu$s | |
| 2 | open | $3 \times 1$ $\mu$s | |
| Brn1 + Ycg1 + DNA (18-bp) | | | |
| 3 | closed | $3 \times 1$ $\mu$s | |
| 4 | open | $3 \times 1$ $\mu$s | |
| **Steered MD Simulation** | | | |
| **System** | **state** | **Duration** | |
| Brn1 + Ycg1 | | | |
| 1 | closed | $3 \times 2500$ ps | |
| Brn1 + Ycg1 + DNA (18-bp) | | | |
| 2 | closed | $3 \times 2500$ ps | |
| **CG MD Simulation** | | | |
| **System** | $\varepsilon_{\mathbf{LB}}$ | $\varepsilon_{\mathbf{HD}}$ | **Duration** |
| Brn1 + Ycg1 | | | |
| 1 | 0.3 | - | $10 \times 10$ $\mu$s |
| 2 | 0.35 | - | $10 \times 10$ $\mu$s |
| 3 | 0.37 | - | $10 \times 10$ $\mu$s |
| 4 | 0.4 | - | $10 \times 10$ $\mu$s |
| 5 | 0.45 | - | $10 \times 10$ $\mu$s |
| 6 | 0.5 | - | $10 \times 10$ $\mu$s |
| Brn1 + Ycg1 + DNA duplex (98-bp) | | | |
| 7 | 0.3 | 0.5 | $50 \times 10$ $\mu$s |
| 8 | 0.3 | 0.6 | $50 \times 10$ $\mu$s |
| 9 | 0.3 | 0.7 | $50 \times 10$ $\mu$s |
| 10 | 0.35 | 0 | $10 \times 10$ $\mu$s |
| 11 | 0.35 | 0.3 | $50 \times 10$ $\mu$s |
| 12 | 0.35 | 0.5 | $50 \times 10$ $\mu$s |
| 13 | 0.35 | 0.6 | $50 \times 10$ $\mu$s |
| 14 | 0.35 | 0.7 | $50 \times 10$ $\mu$s |
| 15 | 0.35 | 0.9 | $50 \times 10$ $\mu$s |
| 16 | 0.35 | 1.0 | $10 \times 10$ $\mu$s |
| 17 | 0.37 | 0.6 | $50 \times 10$ $\mu$s |
| 18 | 0.4 | 0.5 | $50 \times 10$ $\mu$s |
| 19 | 0.4 | 0.6 | $50 \times 10$ $\mu$s |
| 20 | 0.4 | 0.7 | $50 \times 10$ $\mu$s |
| 21 | 0.45 | 0.6 | $50 \times 10$ $\mu$s |
